# A molecular clutch governs stator engagement and flagellar torque

**DOI:** 10.64898/2026.10.14.759253

**Authors:** Wangbiao Guo, Jian Yue, Kai Zhang, Emma Mao, Jiaqi Wang, Jiaqi Guo, Huaxin Yu, Md A. Motaleb, Chunhao Li, Jun Liu

**Author notes:** These authors contributed equally to this work (Wangbiao Guo, Jian Yue, Kai Zhang).

## Abstract

Motile bacteria must continuously adjust their motility to navigate and colonize diverse environments, including mammalian hosts, yet the underlying mechanisms remain poorly understood. Here, using *in situ* single-particle cryo-electron microscopy, we determine near-atomic structures of the flagellar motor in the Lyme disease spirochete *Borrelia burgdorferi* and uncover a novel mechanism for controlling torque generation. We identify a distinct neutral state in which individual torque-generating MotA_5_B_2_ stator complexes remain incorporated in the motor but adopt an extended, tilted conformation disengaged from the rotor. Two conserved flagellar proteins, FliL and FlcA, form a transmembrane scaffold that retains 16 stator complexes, while FapA, a newly identified multidomain flagellar protein, binds the cytoplasmic base of the stator complex and stabilizes its rotor-disengaged conformation. Strikingly, rotor-engaged and rotor-disengaged stators coexist within individual motors, demonstrating that stator retention and torque engagement are separable states. Our findings establish a motor-retained neutral stator state and reveal a molecular-clutch mechanism that independently controls the engagement of individual stator units with the rotor, enabling rapid and precise regulation of flagellar torque and motility.

## Main

Bacterial flagellar motors are remarkable rotary nanomachines that enable cells to navigate complex and rapidly changing environments^1-3^. In model bacteria such as *Escherichia coli* and *Bacillus subtilis*, chemotactic signaling through CheY controls flagellar switching^4,5^, while the second messenger cyclic-di-GMP (c-di-GMP) and its effectors modulate motility in response to surface and physiological cues, promoting transitions between motile and sessile states^6-8^. Stator complexes can also exchange between membrane-diffusing and motor-associated pools, allowing the number of torque-generating units to adapt to mechanical demand^9,10^. These mechanisms regulate motor output through signaling or stator recruitment, but whether torque can be controlled at the level of individual motor-associated stator complexes remains poorly understood.

This question is particularly relevant for pathogenic spirochetes, whose distinctive motility enables movement through complex environments and promotes dissemination and infection. The Lyme disease spirochete *Borrelia burgdorferi* encounters markedly different physical environments as it cycles between *Ixodes* ticks and mammalian hosts and exhibits heterogeneous motility, transitioning among non-motile, wriggling, lunging, and translational states^11-13^. These behaviors are driven by periplasmic flagella and their membrane-embedded motors, in which stator complexes couple ion flow to torque generation and transmission to the rotor^14,15^. Unlike the dynamic stator exchange observed in *E. coli* and *Vibrio* species^9,10^, each spirochetal motor contains 16 stator complexes organized and stabilized by a prominent periplasmic collar^16,17^. This architecture supports the high torque required to rotate periplasmic flagella and deform the cell body but poses a fundamental regulatory question: how can motor output be modulated when stator complexes remain incorporated within the motor?

Here, using *in situ* single-particle cryo-electron microscopy (cryo-EM), we determine near-atomic structures of the intact *B. burgdorferi* flagellar motor in distinct stator-engagement states *in vivo*. Combined with genetic, motility, and cryo-ET analyses, these structures reveal a distinct motor-retained neutral state and a newly recognized molecular clutch that separates stator incorporation from rotor engagement, providing a mechanism for independent control of individual torque-generating units.

## Results

### *In situ* cryo-EM reveals a motor-retained neutral stator state

To visualize stator organization within the native flagellar motor, we applied *in situ* single-particle cryo-EM to intact Δ*cheY3 B. burgdorferi* cells and determined high-resolution structures of their periplasmic flagellar motors *in vivo* (**Fig. 1a**). To overcome the challenge of detecting and aligning motor particles within the crowded cellular environment, we developed an advanced workflow combining manual particle picking with machine-learning-based detection using Topaz^18^, followed by iterative classification and refinement in CryoSPARC^19^. This approach yielded 44,365 motor particles from 24,011 cryo-EM micrographs, enabling 3D reconstruction of the intact flagellar motor, including the stator complexes, MS-ring, rod, collar, C-ring, export apparatus, and ATPase complex (**Fig. 1b-e, supplementary Fig. 1**).

**Fig. 1.**
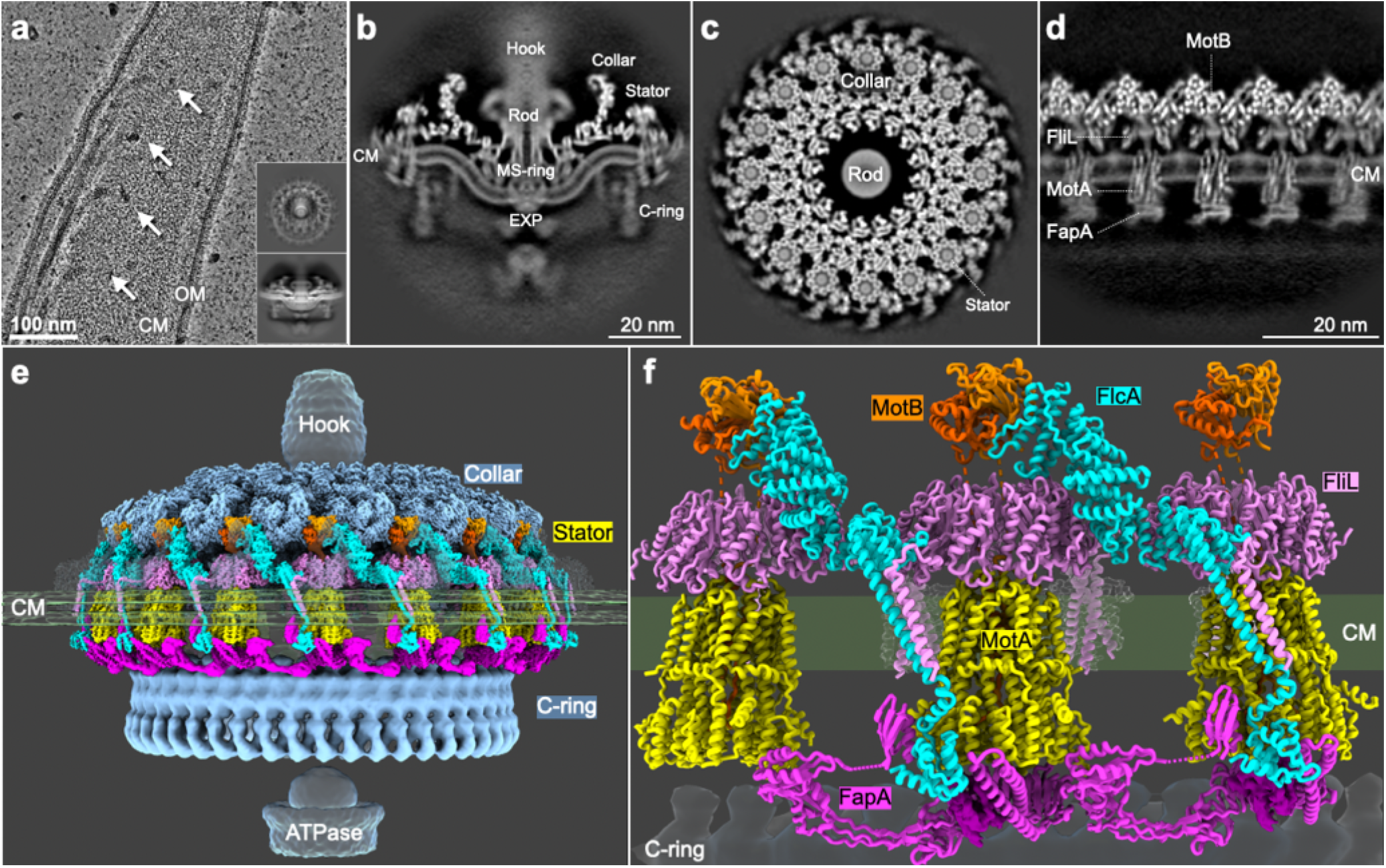
*In situ* cryo-EM structure of the *B. burgdorferi* flagellar motor. (**a**) Representative cryo-EM image of the *B. burgdorferi* cell tip, showing the periplasmic flagellar motors (white arrows), filaments, outer membrane (OM), and cytoplasmic membrane (CM). Representative 2D class averages from side and top views are shown in the bottom insets. (**b**) Cross-sectional side view of the 3D reconstruction of the periplasmic flagellar motor, showing stator complexes that remain anchored within the motor but are disengaged from the C-ring, precluding direct torque transmission. This configuration is designated the neutral state. (**c**) Cross-sectional top view of the 3D reconstruction, with the stator complex, rod, and collar indicated. (**e**) Unrolled side view of the flagellar motor, showing the MotA-MotB stator complexes, together with FliL and FapA. (**e**) Cryo-EM density map of the intact periplasmic flagellar motor, showing the hook, collar, stator complexes, ATPase, and C-ring. (**f**) Enlarged view of the atomic model of the MotA_5_MotB_2_ stator complexes together with the FliL ring, FlcA, and FapA.

Focused classification of the stator complexes and C-ring resolved three distinct motor conformations (**supplementary Fig. 2**). In two conformations, the stator complexes contacted the C-ring in arrangements corresponding to the clockwise (CW) and counterclockwise (CCW) rotational states^20^. The third conformation, however, exhibited a markedly different stator arrangement: the stator complexes remained incorporated within the collar but were displaced outward from the C-ring, eliminating direct stator-rotor contact (**Fig. 1b, supplementary Fig. 2**). Asymmetric reconstruction of individual stator-collar units resolved both the stator complexes and their associated collar components at high resolution, revealing a fixed orientation of the stator complex relative to the collar. Thus, the stator complex remained anchored within the motor but was disengaged from the rotor, precluding direct torque transmission. We therefore designated this configuration a neutral state.

Focused refinement of individual stator complexes in the neutral state together with their associated collar components yielded an *in situ* structure at an overall resolution of 3.79 Å. This near-atomic map enabled modeling of the MotA_5_B_2_ stator complex and surrounding proteins, including FliL ^21,22^ and collar protein FlcA ^23,24^ (**Fig. 1e, f, supplementary Fig. 3, Tables 1-2, movie 1**). On the cytoplasmic side, we identified a previously unrecognized component, FapA, bound beneath the MotA pentamer (**Fig. 1f**). Together, these structures provide a molecular basis for understanding how stator complexes are retained and regulated in the *B. burgdorferi* flagellar motor.

### The neutral stator complex adopts an extended and tilted conformation

The neutral stator complex retained the canonical MotA_5_B_2_ architecture ^25,26^ but adopted an extended configuration tilted by ∼16° relative to the membrane normal (**Fig. 2a-c**). MotB spanned ∼162 Å from its N-terminal transmembrane helix (MotB_TMH_) to its C-terminal peptidoglycan-binding domain (MotB_PGB_) (**Fig. 2c**). Both regions were well resolved, with clear side-chain densities, whereas the intervening region (residues 45-110) was less ordered. Continuous low-resolution density delineated an extended linker passing through the FliL ring, whereas the canonical plug conformation that blocks the MotA channel was not observed in this neutral state (**Fig. 2c**).

**Fig. 2.**
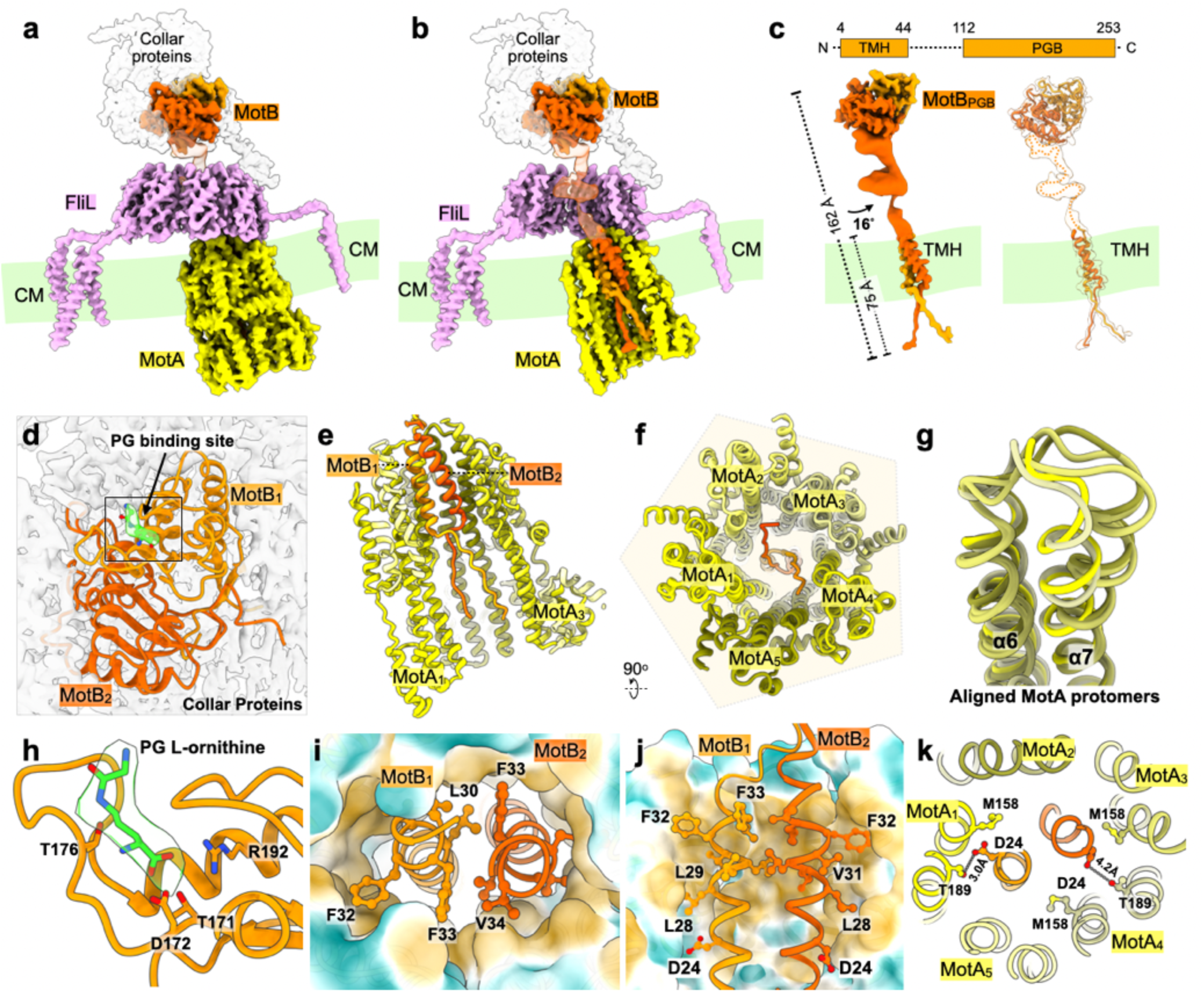
Near-atomic structure of the MotA_5_B_2_ stator complex in a neutral state. (**a**) Cryo-EM density map of the MotA_5_B_2_-FliL complex embedded in the cytoplasmic membrane (CM). Collar proteins interacting with MotB are segmented and shown as transparent white densities. (**b**) Half-section side-view of the MotA_5_B_2_-FliL complex cut along the central axis. MotB dimer is colored as orange and orange-red. (**c**) Cryo-EM density map and atomic model of MotB. The transmembrane helix and the C-terminal peptidoglycan-binding domain are well resolved with clear side-chain densities, whereas the middle region proximal to FliL is poorly resolved. MotB adopts a 16°-tilted, extended conformation relative to the rod axis, spanning ∼162 Å in length, with the transmembrane helix extending ∼75 Å. (**d**) Close-up view of the peptidoglycan-binding (PGB) domain of MotB, showing the peptidoglycan-binding site and its embedding within the collar proteins. (**e**) Atomic model of the MotA_5_B_2_ complex. (**f**) Bottom view of the MotA_5_B_2_ model, highlighting the pentameric organization of MotA. (**g**) Structural alignment of the five MotA protomers, showing their variable loops. (**h**) Cryo-EM density in the PG-binding pocket of MotB, modeled as an L-ornithine (L-Orn) moiety, a characteristic component of the peptide stem in B. burgdorferi peptidoglycan. (**i, j**) Top and side views of the stator complex, showing the MotB transmembrane helices enclosed within a hydrophobic environment formed by the MotA pentamer, with no apparent continuous solvent-accessible channel. (**k**) Close-up view of the ion-binding sites, highlighting their distinct local interaction environments. The distance between the conserved Asp24 of MotB_1_ and Thr189 of MotA_1_ is ∼3.0 Å, forming a hydrogen bond, whereas the corresponding distance in MotB_2_ is ∼4.2 Å.

The MotA pentamer exhibited a nearly symmetric organization. Its five protomers adopted similar conformations, superimposing with an RMSD of less than 1 Å and forming an approximately pentagonal assembly when viewed along the central axis (**Fig. 2f, g**). Conformational differences were largely confined to flexible periplasmic loops that interact with MotB and the surrounding FliL ring (**supplementary Fig. 4**). This near-symmetric organization contrasts with the asymmetric MotA_5_B_2_ conformations observed in previously characterized stator structures determined *in vitro*^*25,26*^.

The MotB_PGB_ domains were largely buried within the surrounding collar proteins, while their peptidoglycan-binding surfaces remained exposed. Additional density within the conserved peptidoglycan-binding pocket was modeled as an L-ornithine (L-Orn) moiety, a characteristic component of the peptide stem in *B. burgdorferi* peptidoglycan ^27^ (**Fig. 2d-h, supplementary Fig. 4**). These observations suggest that MotB anchors the neutral stator complex through interactions with both the collar and the peptidoglycan layer.

Despite the absence of resolvable density for the canonical MotB plug, the ion-conduction pathway appeared occluded. The two MotB_TMH_ helices were enclosed within the MotA pentamer and surrounded by closely packed hydrophobic side chains, leaving no apparent continuous solvent-accessible channel (**Fig. 2i-j)**. This hydrophobic packing may restrict ion conduction and thereby limit unproductive ion flux when the stator complex is disengaged from the rotor.

The *in situ* structure further distinguished the two MotB subunits based on their positions relative to the rotor, designated rotor-proximal MotB_1_ and rotor-distal MotB_2_ (**Fig. 2k**). In MotB_1_, the conserved ion-binding residue Asp24 lay 3.0 Å from Thr189 of MotA_1_, consistent with a potential hydrogen-bonding interaction. By contrast, the corresponding Asp24-Thr189 distance in MotB_2_ was 4.2 Å (**Fig. 2k**). Thus, despite the near-symmetric organization of the MotA pentamer, the two MotB ion-binding sites retained distinct local interaction environments.

Collectively, these features define a distinctive neutral stator architecture characterized by an extended, tilted conformation and an occluded ion-conduction pathway.

### FliL forms interconnected transmembrane nonamers that organize stator complexes

FliL has been implicated in stator retention and enhanced motility^21,22^. Our *in situ* structure revealed that FliL forms a membrane-anchored nonameric ring surrounding the periplasmic portion of the MotA_5_B_2_ stator complex (**Fig. 3a-c**). In the neutral state, the periplasmic surface of MotA contacts the inner wall of the FliL ring through interactions involving MotA_3_ and MotA_4_ and the collar-facing FliL_6_ and FliL_7_ subunits. These contacts may constrain MotA within the FliL ring and stabilize its rotor-disengaged position. Each FliL protomer comprises a transmembrane helix and a long linker (residues 1-77) connecting it to the periplasmic domain (**Fig. 3d**). In contrast to the ordered periplasmic ring, the transmembrane helices splayed radially and exhibited pronounced structural heterogeneity (**Fig. 3b-d**). Only four of the nine helices, belonging to FliL_1_, FliL_2_, FliL_5_, and FliL_9_, were sufficiently resolved for nearly complete modeling. The transmembrane helices of FliL_1_, FliL_2_, and FliL_9_ formed bundles with neighboring membrane-associated collar components, including FlbB, whereas the FliL_5_ helix contacted FlcA (**Fig. 4b**). These interactions collectively anchor the FliL ring within the membrane-associated collar.

**Fig. 3.**
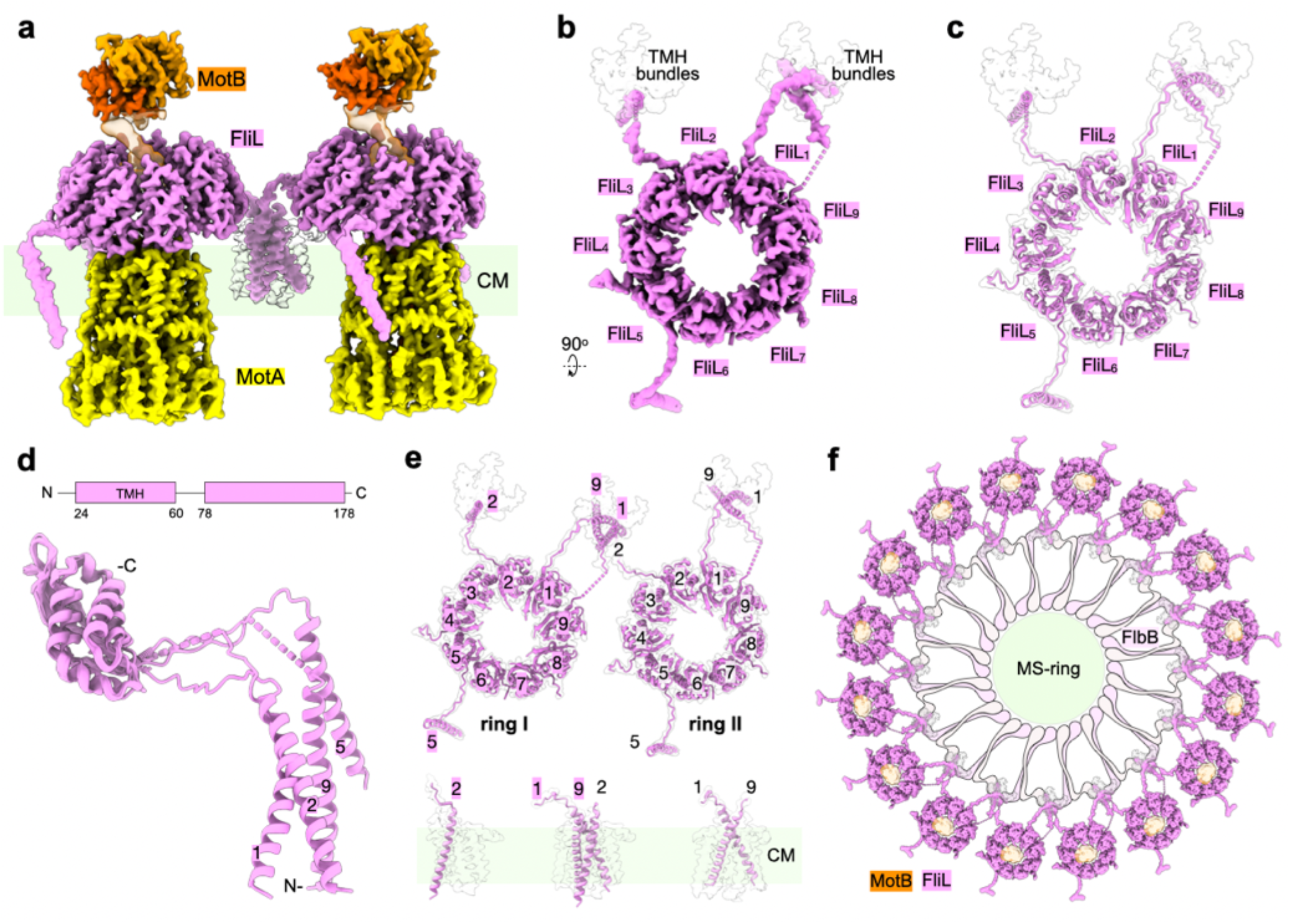
Structure of the transmembrane FliL nonamer. (**a**) Cryo-EM density map of two adjacent MotA_5_B_2_- FliL supercomplexes. (**b**) Cryo-EM density map of the FliL nonamer. Four FliL protomers contain well-resolved transmembrane helices (TMHs). On the inner side of the FliL ring facing the basal body, a bundle of seven TMH densities is observed, three of which are assigned to FliL. (**c**) Near-atomic model of the FliL nonamer. FliL_1_, FliL_2_, FliL_5_, and FliL_9_ were modeled with nearly complete TMHs. (**d**) Structural superposition of FliL monomers. (**e**) Two adjacent FliL nonamers (top) showing inter-ring interactions, in which FliL_1_ and FliL_9_ from one nonamer interact with FliL_2_ of the neighboring nonamer. The close-up view (bottom) shows interactions of FliL_1_, FliL_2_, and FliL_9_ with transmembrane helices of collar proteins (shown as transparent withe). (**f**) Arrangement of the sixteen FliL nonamers surrounding the motor.

**Fig. 4.**
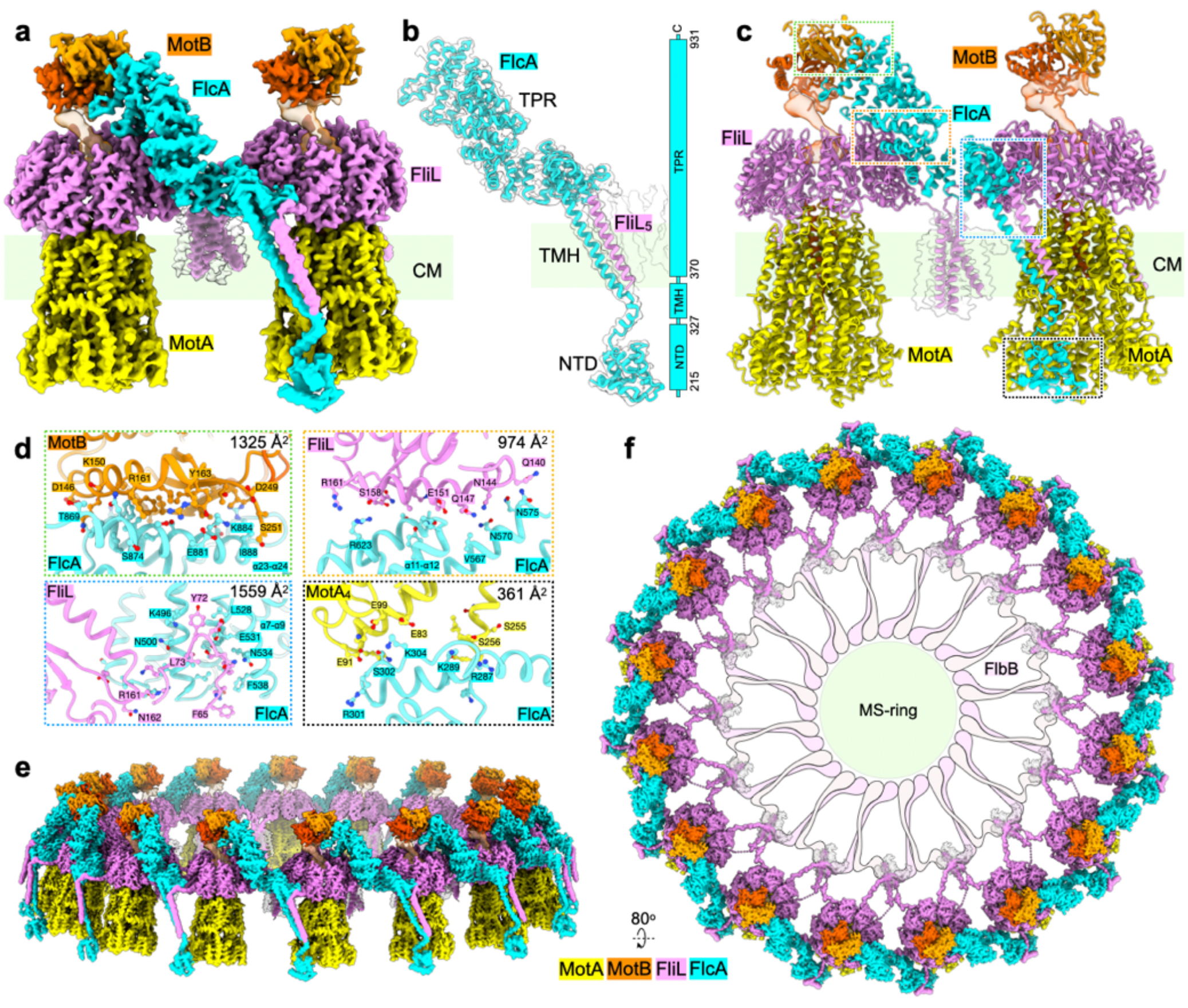
FlcA bridges MotA and the neighboring MotB. (**a**) Cryo-EM density maps of two adjacent MotA_5_B_2_- FliL complexes connected by one FlcA monomer. (**b**) Near-atomic model of FlcA with the corresponding cryo-EM density map shown in transparent white. FlcA interacts with FliL_5_ through their transmembrane helices within the cytoplasmic membrane (CM). FlcA comprises a cytoplasmic globular N-terminal domain, a single transmembrane helix (TMH), and a periplasmic tetratricopeptide repeat (TPR) domain. (**c**) Near-atomic model of two adjacent stator-FliL complexes connected by one FlcA monomer. (**d**) Close-up views of the interaction interfaces between FlcA α23-α24 and MotB_1_ (green box), FlcA α5-α7 and FliL_5_ (orange box), FlcA α12-α15 and FliL_8_ (dark blue box), and the FlcA N-terminal domain and MotA_4_ (black box). (**e**) Oblique view of the cryo-EM density map of the 16 FlcA-FliL-MotAB supercomplex. (**f**) Top view of the 16 FlcA-FliL-MotAB supercomplexes, with FlbB and the MS-ring shown schematically at the basal body.

Notably, the transmembrane helices also interconnected adjacent FliL rings. FliL_1_ and FliL_9_ from one nonamer contacted FliL_2_ from the neighboring ring, forming interlocking transmembrane helix bundles (**Fig. 3e**). These interactions connect the 16 FliL rings into a continuous circumferential scaffold surrounding the rotor (**Fig. 3f**).

### FlcA forms a transmembrane bridge linking neighboring MotAB-FliL complexes

FlcA is a component of the flagellar collar in *B. burgdorferi*, and its loss causes defects in motor assembly and flagellar motility (**Fig. 4a, supplementary Table 3**)^23,24,28^. Our *in situ* structure revealed how FlcA integrates neighboring stator complexes through interactions on both sides of the cytoplasmic membrane. FlcA comprises a globular N-terminal cytoplasmic domain (NTD), a single transmembrane helix, and an elongated periplasmic tetratricopeptide repeat (TPR) domain containing 24 α-helices (**Fig. 4b**).

On the periplasmic side, the TPR domain forms a superhelical scaffold that bridges two adjacent FliL nonamers. Its interfaces with FliL_5_ of one ring and FliL_8_ of the neighboring ring bury 1,559 Å^2^ and 974 Å^2^ of surface area, respectively (**Fig. 4c-d**). At the distal end of the TPR domain, helices α23-α24 contact the peptidoglycan-binding domain of MotB through an interface burying 1,325 Å^2^, providing a major interaction site for MotB_PGB_ within the collar (**Fig. 4c-d**). On the cytoplasmic side, FlcA_NTD_ contacts the outward-facing surface of MotA_4_ through an interface burying 361 Å^2^ (**Fig. 4c-d**). Thus, a single FlcA molecule bridges neighboring FliL rings while simultaneously linking periplasmic MotB to cytoplasmic MotA across the membrane. This arrangement is repeated around the motor, with 16 FlcA molecules interconnecting the MotAB-FliL assemblies into an integrated circumferential stator array (**Fig. 4c-f**).

### FapA masks the stator-rotor interface in the neutral state

On the cytoplasmic side, our *in situ* reconstruction revealed FapA bound to the base of the MotA pentamer at its torque-generating surface (**Fig. 5a-b**). This position places FapA directly at the stator-rotor interface, suggesting a role in regulating MotA engagement with the C-ring.

**Fig. 5.**
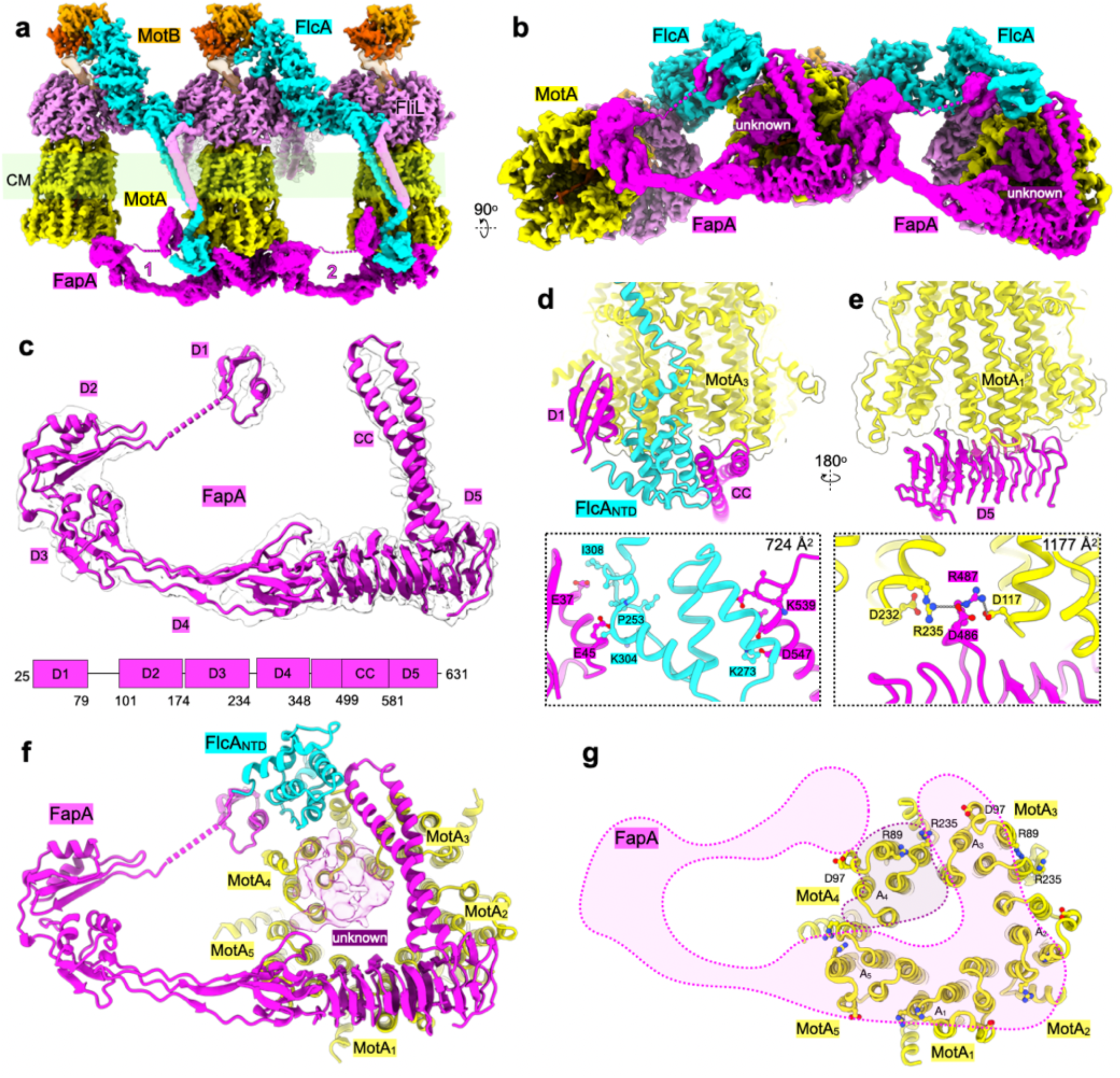
FapA occludes the MotA interface. (**a**) Cryo-EM density map of three adjacent FliL-MotAB supercomplexes together with two copies of FapA, and FlcA. (**b**) Bottom view of the corresponding density map, showing FapA completely occluding the MotA-C-ring interaction interface, with a small, unassigned density between FapA and MotA. (**c**) Structural model of an individual FapA monomer. FapA comprises five domains, as indicated. (**d, e**) Two close-up views of the interactions between the D1 and D5 domains of FapA and the FlcA N-terminal domain and MotA. (**f**) Bottom view of FapA and unknown density in the context of the MotA pentamer. (**g**) FapA and unknown density occupies the cytoplasmic surface of MotA, occluding the MotA-C-ring interaction interface. The conserved C-ring-interacting residues Asp97, Arg89, and Arg235 of MotA are mostly blocked by FapA and unknown density (outlined by dash lines).

FapA formed an elongated structure comprising five domains, D1-D5 (**Fig. 5c-d**). D1-D3 adopted compact α+β ferredoxin-like folds, whereas D4 formed a β-sandwich-like fold. The C-terminal D5 domain comprised a triangular β-solenoid followed by an extended two-helix coiled coil. D1-D4 were resolved at 6-8 Å resolution, allowing their overall secondary-structure organization to be discerned, while D5 was resolved at ∼3.5 Å with well-defined side-chain densities (**Fig. 5c-d**). Comparative sequence and structural analyses identified FapA homologues in diverse flagellated bacteria, including species outside the spirochetes, with the D5 domain showing the greatest conservation (**supplementary Fig. 5**).

FapA_D5_ formed the principal MotA-binding module (**Fig. 5e-g**). Its triangular β-solenoid contacted the rotor-facing MotA_1_, MotA_2_, and MotA_5_ subunits through an interface burying approximately 1,777 Å^2^. A loop comprising residues 484-491 inserted into the cleft between MotA_1_ and MotA_5_, occupying the region containing charged residues Asp117, Asp232, and Arg235 implicated in FliG binding at the C-ring (**Fig. 5f-g**). The extended coiled coil traversed the base of the pentamer to contact MotA_3_ and MotA_4_ (**Fig. 5f, j**). Thus, FapA_D5_ contacted all five MotA subunits and masked a substantial portion of the rotor-facing surface, providing a structural basis for preventing productive stator**-**rotor engagement in the neutral state.

FapA also linked the MotA pentamer to the collar scaffold. FapA_D1_ contacted FlcA_NTD_ through an interface burying 724 Å^2^, while the central D2-D4 domains extended laterally toward a neighboring MotA complex (**Fig. 5a, d and e**). This arrangement anchors FapA-bound MotA to FlcA and positions FapA to form connections between adjacent stator assemblies on the cytoplasmic side. Notably, the FapA density was not resolved beneath CW- or CCW-engaged stator complexes, in which MotA contacted the C-ring (**supplementary Fig. 2**). Together, these structures support a molecular-clutch model in which FapA masks the MotA-FliG interface in the neutral state; its dissociation or repositioning may allow rotor engagement.

### FapA supports stator organization and flagellar motility

To understand the functional role of FapA, we compared the motility of wild-type, Δ*fapA*, and complemented (*C*Δ*fapA*) strains. Δ*fapA* cells remained motile but exhibited impaired swimming compared with wild-type and complemented cells (**Fig. 6a-b**). Swimming-plate assays showed a modest but significant reduction in colony expansion, and single cell tracking in 1% methylcellulose revealed a significant decrease in swimming velocity. Importantly, Δ*fapA* cells exhibited a pronounced defect in directional reversal. Wild-type cells reversed frequently (∼25 times per minute), whereas Δ*fapA* cells reversed substantially less frequently (∼7 times per minute), resulting in more persistent movement in one direction (**Fig. 6a-c, supplementary Fig. 6, movies 2-3**).

**Fig. 6.**
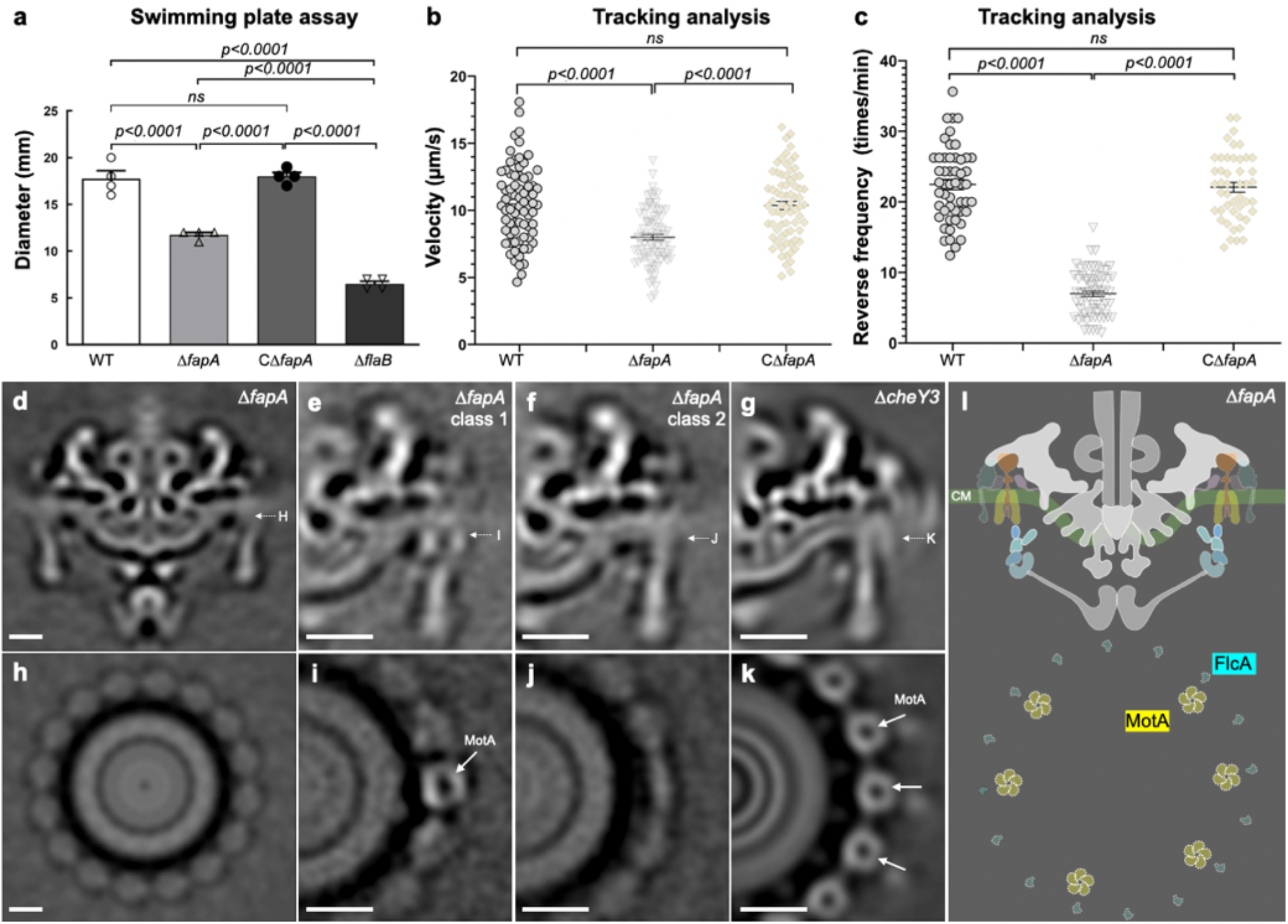
Characterization of FapA and its impacts on stator assembly and flagellar motility. (**a**) Swimming motility assay on the plates of wild-type (WT), Δ*fapA*, and complemented (*C*Δ*fapA*) strains. (**b**) Single-cell tracking analysis showing swimming velocity of WT, Δ*fapA* and *C*Δ*fapA* strains. (**c**) Directional reversal of WT, Δ*fapA* and *C*Δ*fapA* strains. (**d**-**f, h-j**) Structural analyses of Δ*fapA* motors by cryo-ET and subtomogram averaging. Stator-associated densities are not well resolved in the global average (d, h). Classification reveals that only ∼33% of stator complexes are detectable (e, i), whereas the remaining stator complexes are not resolved (f, j). (**g, k**) *In situ* structures of Δ*cheY3* motors in a counterclockwise (CCW) state, shown as controls, in which the stator complexes are clearly resolved. (**l**) Cartoon model of the Δ*fapA* motor illustrating increased stator dynamics.

To investigate the molecular basis of these motility defects, we examined Δ*fapA* motors by cryo-electron tomography (cryo-ET) and subtomogram averaging (**Fig. 6d-g, h-j, and supplementary Fig. 7**). In contrast to Δ*cheY3* motors (**Fig. 6h, k**), Δ*fapA* motors lacked an ordered density corresponding to MotAB-FliL-FlcA (**Fig. 6d-g**). Only ∼33% of stator complexes were resolved through focused classification (**Fig. 6e, i**). Thus, loss of FapA markedly disrupts the organization of stator-associated components without abolishing motility. Together with the reductions in swimming velocity and reversal frequency, these findings demonstrate that FapA contributes to stable organization of the stator-collar assembly and supports efficient flagellar motility and its regulation.

## Discussion

For decades, flagellar stator complexes were conventionally thought to be torque-generating units statically anchored within the motor^29^. Single-molecule studies subsequently established that stator complexes dynamically exchange between motor-bound and membrane-diffusing pools^9,10^, while recent studies suggested that ion flow drives rotational motion of the MotA pentamer around MotB during torque generation^20,25,26,30^. Our *in situ* structures of the *B. burgdorferi* flagellar motor reveal a neutral state in which the MotA_5_B_2_ stator complex remains incorporated within the motor but is disengaged from the rotor, while its ion-conduction pathway is occluded (**Fig. 1**). Thus, the neutral state defines a previously unrecognized configuration: motor associated yet functionally disengaged. Importantly, our structures further show that individual stator units within the same motor can transition between torque-generating and rotor-disengaged configurations without dissociating from the motor (**Fig. 7a-h, and supplementary Video 4**), revealing an additional layer of stator dynamics and regulation beyond exchange between motor-bound and membrane-diffusing pools.

**Fig. 7.**
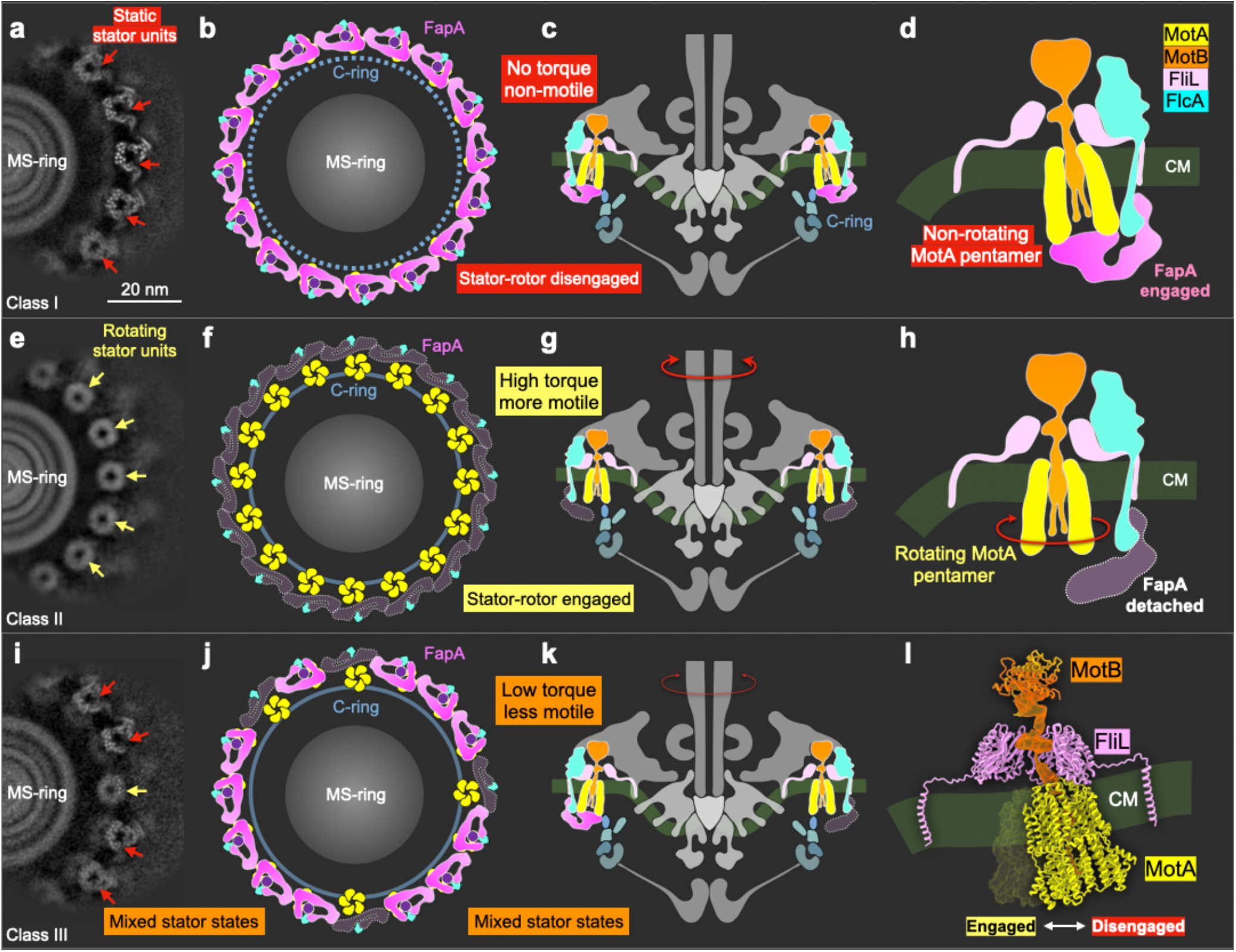
A molecular-clutch model for regulation of flagellar torque. (**a-d**) Neutral state (EMD-79275). FliL and FlcA retain MotA_5_B_2_ within the motor, whereas FapA forms a molecular clutch that stabilizes the rotor-disengaged stator configuration. The stator complex remains anchored to the FliL-FlcA-FapA scaffold but is disengaged from the C-ring. (**e-h**) High-torque rotating state (EMD-79272). FapA is released from all stator complexes, allowing MotA_5_B_2_ to engage with the C-ring and generate torque for rotor rotation. (**i-l**) Low-torque rotating state (EMD-79274). Only a subset of stator complexes is engaged with the C-ring, resulting in reduced torque generation. FapA-mediated transitions between rotor-disengaged and rotor-engaged stator configurations provide a mechanism for regulating individual stators, thereby controlling the number of torque-generating units and modulating motor output. The stator complex is constrained within the FliL ring, with MotA adopting distinct conformations in the engaged and disengaged states.

The neutral state is stabilized by an interconnected transmembrane scaffold and a cytoplasmic regulatory complex. FliL encircles individual stator complexes, whereas FlcA bridges neighboring FliL rings and connects MotB in the periplasm to MotA on the cytoplasmic side (**Fig. 4**). Together, FliL and FlcA retain 16 stator complexes within the motor while preserving the flexibility required for individual stator regulation. Anchored to the scaffold through FlcA, FapA extends across the rotor-facing surface of MotA and masks residues required for productive interaction with the C-ring. The stator complex is therefore both spatially displaced from the C-ring and sterically prevented from establishing a torque-generating interface. Critically, the coexistence of rotor-engaged and rotor-disengaged stator units within the same motor provides a molecular basis for independent stator control, allowing neighboring units to remain engaged and continue transmitting torque (**Fig. 7**). Consistent with this model, Δ*fapA* motors remain assembled but lose ordered stator-associated density, while mutant cells exhibit reduced swimming velocity and reversal frequency (**Fig. 6**). Thus, FapA acts as an integrated molecular clutch that contributes to the stability of motor-associated stator complexes and control their engagement with the rotor.

Such a mechanism may be particularly advantageous for the high-torque motors of spirochetes, in which numerous stator complexes are embedded within an elaborate periplasmic collar^28,31,32^, as well as other high-torque polar motors, including those of *Helicobacter pylori* ^33,34^ and *Campylobacter jejuni* ^35-37^ **(Supplementary Fig. 8)**. By modulating the number of engaged stator complexes without requiring their release from the motor, this mechanism may preserve the structural integrity of the stator array while enabling rapid and graded adjustment of motor output. The pronounced reduction in reversal frequency in Δ*fapA* cells raises the possibility that such regulation contributes to transitions between motility states, although how this process is controlled remains unknown. Intriguingly, we also observed an unassigned globular density and a small ring-shaped density adjacent to FapA that may represent additional regulatory components coupling upstream signals to stator engagement. Their identities and functions remain to be determined.

In summary, our findings support an integrated molecular-clutch model in which individual stator complexes can disengage from the rotor while remaining incorporated within the flagellar motor. By separating stator retention from rotor engagement, this mechanism enables the number of torque-generating units to be adjusted without disrupting the overall stator array, providing a potential means for rapid and precise control of motor output as bacteria navigate and colonize diverse environments. The conservation of FapA and associated components beyond spirochetes suggests that this mechanism extends to diverse bacterial flagellar motors.

## Methods

### *In situ* cryo-EM/ET sample preparation

Δ*cheY3 B. burgdorferi* cells ^20^ were pelleted by centrifugation at 10,000*g* for ∼10 min, gently washed with phosphate-buffered saline (PBS), and resuspended in PBS to an optical density (OD_600_) of 1.0. Cryo-EM grids (200 mesh R2/1 Quantifoil copper grids or 200 mesh Lacey carbon copper grids, Ted Pella, Inc) were glow-discharged for ∼30 s before use. A 5 μL aliquot of the cell suspension was applied to each grid, blotted from the front with filter paper (Whatman™) for ∼8 s, and plunge-frozen into a liquid ethane-propane mixture using a GP2 plunge freezer (Leica). The environmental chamber was maintained at 25 °C and 90% relative humidity throughout vitrification.

### Cryo-EM data collection and processing

Cryo-EM specimen was imaged using two 300 kV Titan Krios electron microscopes (Thermo Fisher Scientific) for multiple sessions. One equipped with a field emission gun and a post-GIF K3 Summit direct electron detector (Gatan). Images were recorded in nanoprobe mode with the following settings: magnification at 81,000×, spot size at 5, illumination area at 1.22 µm, slit width at 20 eV, C2 aperture at 100 µm, and objective aperture at 100 µm. A customized multishot script was used to directly image the cell tips on the cryo-EM grids with SerialEM^38^, applying defocus values from **-**1.6 µm to **-**2.2 µm. The total electron dose was ∼70 e^-^/Å^2^. Movies were recorded in super-resolution mode with a nominal pixel size of 0.534 Å/pixel and motion-corrected with 2× binning to an effective pixel size of 1.068 Å/pixel. Another Titan Krios was imaged with similar setting, except at an illumination area of 1.05 µm. Movies were recorded with a nominal pixel size of 0.5311 Å/pixel and motion-corrected with 1× binning and scale to an effective pixel size of 1.068 Å/pixel. Collectively, a total of 24,011 micrographs were imaged from multiple sessions.

All micrographs were motion-corrected ^39^ and subjected to Patch CTF Estimation in cryoSPARC^19^. An initial set of ∼5,000 flagellar motors were manually picked. Particles were extracted with a box size of 900 pixels and Fourier-cropped to 512 pixels. An initial model was generated by *ab initio* reconstruction using the manually picked particles with C16 symmetry. Around 1,000 particles exhibiting well-defined collar densities were selected for initial reference. These particles were used to train the machine-learning particle picker Topaz^40^, which was subsequently applied to all micrographs. In total, 44,365 periplasmic flagellar motor particles were selected.

After including all particles, homogeneous refinement and non-uniform refinement with C16 symmetry yielded a reconstruction at 7.05 Å resolution (FSC = 0.143). To resolve conformational heterogeneity of stator-C-ring interface, focused 3D classification was performed masking on the collar, stator, and C-ring regions, respectively. The masks were generated from the resolved density maps. Based on the conformations of the stator and C-ring, particles were classified into CW, CCW, and neutral states. Local refinement with C16 symmetry-expansion particles yielded reconstructions of the CW state at 6.41 Å (178,499 particles), the CCW state at 7.76 Å (40,884 particles), and the neutral state at 4.75 Å (490,115 particles).

To improve the local resolution of the stator complex, all particles were recentered to one quarter of the motor, re-extracted with a box size of 640 pixels, and Fourier-cropped to 448 pixels. Following homogeneous refinement, local CTF refinement, and local refinement with C1 symmetry, the overall resolution improved to 3.72 Å. Focused 3D classification of the stator region was then performed to separate the CW, CCW, and neutral conformations. New masks were generated from the improved density maps, particularly for the FapA architecture. The final local refinements yielded reconstructions of the neutral state at 3.53 Å (425,097 particles). The cryo-EM maps were further processed using the deep learning-based refinement algorithm DeepEMhancer ^41^ or EMReady ^42^ to enhance visualization.

### Bacterial strains and growth conditions

Low-passage *B. burgdorferi* sensu stricto strain B31-A3 A3-68 ^43^ and its derivative mutants were cultured in Barbour-Stoenner-Kelly II (BSK-II) liquid medium or on semi-solid agar plates at 34°C under 3.4% CO_2_ as described previously^44,45^. Selective antibiotics were added to the media as needed: kanamycin (300 μg/ml) or gentamicin (40 μg/ml). *Escherichia coli* NEB 5α strain (New England Biolabs, Ipswich, MA, USA) was utilized for plasmid cloning and propagation. Antibiotics selection for *E. coli* transformants was maintained using gentamicin (5 μg/ml), or ampicillin (100 μg/ml).

### Construction of the *fapA* (*BB0267*) in-frame deletion mutant and its isogenic complemented strain

To construct an in-frame deletion of *fapA* (gene locus, *BB0267*), targeted gene disruption vectors were engineered via overlap extension PCR as described previously^22^. Briefly, primer pairs P_3_/P_4_ and P_5_/P_6_- containing sequence overlaps complementary to the kanamycin resistance cassette (*aph1*) were designed to amplify approximate 1-kb regions upstream and downstream of the *fapA* coding sequence. Following initial amplification of the individual 5’- and 3’-flanking and *aph1* fragments, a fusion PCR was performed using primers P_3_/P_6_ to assemble all three fragments into the final *fapA::aph1*disruption construct. Mutant clones were selected on semi-solid BSK-II agar plates supplemented with kanamycin (300 μg/ml).

To generate a *trans*-complementation construct for the *fapA* mutant, the *B. burgdorferi flgB* promoter (*P*_*flgB*_) was PCR-amplified using primers P_7_/P_8_ containing 5′-BamHI and 3′-overlapping sequence complementary to the *fapA* coding sequence. The intact *fapA* gene was subsequently amplified using primers P_9_/P_10_ engineered with 3′-PstI sites. Two fragments were assembled via fusion PCR using primers P_7_/P_10_. The cassette containing *P*_*flgB*_*-fapA* was ligated to pJET1.2/blunt cloning vector (Thermo Scientific, Waltham, MA), excised using BamHI and PstI, and then subcloned into corresponding sites in shuttle vector pBSV2G to produce the final complementation plasmid, *P*_*flgB*_*-fapA*/pBSV2G. The resulting vector was then transformed into the *fapA* mutant (Δ*fapA*) cells via electroporation, and transformants were selected on semi-solid BSK-II agar plates supplemented with kanamycin (300 μg/ml) and gentamicin (40 μg/ml). The resulting complemented clones were confirmed by PCR. One of positive clones (*C*Δ*fapA*) was selected for further phenotypical analyses. All oligonucleotide primers utilized are listed in **Supplementary Table 4**.

### Gel electrophoresis and immunoblot analysis

Sodium dodecyl sulfate-polyacrylamide gel electrophoresis (SDS-PAGE) and enhanced chemiluminescence (ECL) immunoblotting were performed as described previously^46^. Briefly, whole-cell lysates (10-20 μg protein per lane) were resolved on 12%SDS-PAGE gels and transferred onto polyvinylidene difluoride (PVDF) membrane (Bio-Rad Laboratories, Hercules, CA, USA). Membranes were blocked and incubated with specific primary antibodies, including FlaB monoclonal antibody (H9724, provided by A. G. Barbour, University of California, Irvine, CA), DnaK monoclonal antibody (J. Benach, SUNY at Stony Brook, NY), and polyclonal antibodies against FlgG, FliF, FlgE, FlhF, FliD which were reported in our previous study^47,48^. Blots were incubated with horseradish peroxidase (HRP)-conjugated secondary antibodies and developed using an ECL luminol substrate. Chemiluminescent signals were acquired using a ChemiDoc MP imaging system and quantified with Image Lab software (Bio-Rad Laboratories). All immunoblot analyses were performed using three independent biological replicates.

### Bacterial motion tracking analysis and swimming plate assays

Swimming velocity of *B. burgdorferi* cells was evaluated using a computer-based motion tracking system as described previously^49^. Briefly, logarithmic-phase cultures were diluted 1:1 in BSKII medium, and a 20 μl aliquot was mixed with an equal volume of 2% methylcellulose (4,000 cP viscosity; MC4000). Cell movement was recorded using iMovie software on a Macintosh computer. Videos were exported as QuickTime files and imported into OpenLab (Improvision Inc., Coventry, UK) for frame cropping, calibration, and conversion to LIFF format. Individual cell trajectories and velocities were quantified using Volocity software (Improvision Inc., Coventry, UK). For each strain, at least 20 individual cells were tracked for up to 30 sec. Swimming plate assays was performed using 0.35% agarose plates prepared with BSK-II medium diluted 1:10 in Dulbecco’s phosphate-buffered saline (dPBS, pH 7.5) as previously described^44,50,51^. Wild-type (WT), the *fapA* mutant (Δ*fapA*), and its isogenic complementary strain (*C*Δ*fapA*) were inoculated into the semi-solid agar plate; the non-motile *flaB* mutant served as a negative control to define the initial inoculum size. Plates were incubated for 4 to 5 days at 34°C under 3.4% CO_2_. Swimming ring diameters were measured in millimeters (mm), and mean diameters average values were calculated from four independent biological replicates.

### Cryo-ET and subtomogram averaging

Frozen-hydrated specimens of Δ*fapA B. burgdorferi* cells were imaged on a 200 kV Glacios transmission electron microscope (Thermo Fisher Scientific) equipped with a field emission gun and a K3 Summit direct electron detector (Gatan). Tilt series were collected using SerialEM ^38^ with the FastTomo script ^52^ using a dose-symmetric acquisition scheme. Data were acquired at a defocus of **-**4.8 μm with a cumulative electron dose of ∼70 e^-^/Å^2^ distributed over 33 tilt images spanning **-**48° to +48° at 3° increments. Movie frames were motion-corrected using MotionCor2^39^, followed by tilt-series alignment and stacking in IMOD^53^. Tomograms were reconstructed from 8× binned tilt series using Tomo3D^54^. In total, 123 tilt series were collected from which 769 flagellar motors were manually picked using Tomopick^55,56^. Initial subtomograms were conducted from the 8× binned tomograms with the i3 software package^55,56^, followed by iterative alignment and classification to refine particle positions and remove low-quality particles. Final subtomograms were extracted from the unbinned tomograms using the refined coordinates, and subtomogram averaging was subsequently performed at bin 4.

### Molecular Modeling

Amino acid sequences of FliL (Q57442), MotA (Q44902), MotB (H7C7P3), FlcA (O51305) and FapA (H7C7R5 from *B. burgdorferi* strain B31 (ATCC 35210/DSM 4680/CIP 102532) were retrieved from UniProt and used as input for AlphaFold3 ^57^ to generate predicted monomeric structures **(Supplementary Fig. 9, 10)**. The predicted models were fitted into the cryo-EM map using ChimeraX_58_ followed by interactive manual adjustment in Coot ^59^ and real-space refinement in Phenix^60^. Final models exhibited high stereochemical quality as assessed by Phenix validation tools. Sequence alignments were performed using Clustal Omega^61^. ChimeraX ^58^ was used for cryo-EM map segmentation, visualization, and molecular surface rendering and animation.

## Acknowledgements

We thank Jennifer Aronson for critical reading and editing of the manuscript. We thank Jianfeng Lin, Kangkang Song, Zephan Melville, Jing Wang and Edward Eng for helping on cryo-EM data collection. We thank Dongjie Zhu, Kexin Zhang, and Rui Zhang for advice on structure improvement of the stator complex. We thank Charles Sindelar from Yale Center for Research Computing for providing support on cryo-EM data processing.

## Funding

W.G., J.Y., M.A.M., and J.L. were partly supported by grants R01AI087946 and R01AI132818 from National Institute of Allergy and Infectious Diseases (NIAID) and National Institutes of Health (NIH).K.Z. and C.L were supported by grants R01AI078958 (NIAID), R01DE023080 (NIDCR), and R01DE034063 (NIDCR). Cryo-EM data were collected at the Yale CryoEM Resource funded in part by the NIH grant 1S10OD023603-01A1. Some of data were performed at the National Center for CryoEM Access and Training (NCCAT) and the Simons Electron Microscopy Center located at the New York Structural Biology Center, supported by NIH (Common Fund U24GM129539, NIGMS R24GM154192), the Simons Foundation (SF349247), and NY State Assembly.

## Author contributions

W.G., J.Y., C.L. and J.L. conceived the project. W.G. performed cryo-EM and cryo-ET data collection, image processing, and manuscript writing. J.Y. conducted model building, data analysis, and manuscript writing. K.Z. and C.L. generated the strain and related mutant strains, and performed motility assays, western blots and growth inhibition assays. E.M., J.G., J.W. and H.Y. assisted with data acquisition and particle picking. C.L. and J.L. provided resources, supervision, and technical expertise. W.G., J.Y., C.L. and J.L. analyzed and interpreted the data and wrote the manuscript with contributions from all authors. All authors reviewed and approved the final manuscript.

## Competing interests

The authors declare no competing interests.

## Data availability

The atomic coordinates and corresponding density maps have been deposited in the Protein Data Bank (PDB) and Electron Microscopy Data Bank (EMDB), respectively. The focused-refined structures of the flagellar motor with a single stator unit, two adjacent stator units connected by one FapA, and the overall motor with 16 stator units have been deposited as PDB 38VM/EMD-79182, 38VW/EMD-79187, and 38WF/EMD-79198, respectively (**Supplementary Table 1**). The 3D classification maps representing different stator conformations were reconstructed using homogeneous reconstruction only function. The overall 16-stator motor maps in the neutral, clockwise (CW), and counterclockwise (CCW) states have been deposited as EMD-79260, EMD-79268, and EMD-79270, respectively. Focused classification maps of one-quarter of the motor, representing CW, neutral, and mixed CW/neutral stator states, have been deposited as EMD-79272, EMD-79273, EMD-79274, and EMD-79275 (**Supplementary Table 2**). Source data are provided with this paper.

## Notes

### Competing Interest Statement

The authors have declared no competing interest.

## References

1 Einenkel, R. et al. Building the bacterial flagellum: coordinating regulation, dynamic assembly, and function. Microbiol Mol Biol Rev 89, e0009222 (2025). 10.1128/mmbr.00092-22

2 Wadhwa, N. & Berg, H. C. Bacterial motility: machinery and mechanisms. Nat Rev Microbiol 20, 161–173 (2022). 10.1038/s41579-021-00626-4

3 Chaban, B., Hughes, H. V. & Beeby, M. The flagellum in bacterial pathogens: For motility and a whole lot more. Semin Cell Dev Biol 46, 91–103 (2015). 10.1016/j.semcdb.2015.10.032

4 Kearns, D. B. A field guide to bacterial swarming motility. Nat Rev Microbiol 8, 634–644 (2010). 10.1038/nrmicro2405

5 Porter, S. L., Wadhams, G. H. & Armitage, J. P. Signal processing in complex chemotaxis pathways. Nat Rev Microbiol 9, 153–165 (2011). 10.1038/nrmicro2505

6 Boehm, A. et al. Second messenger-mediated adjustment of bacterial swimming velocity. Cell 141, 107–116 (2010). 10.1016/j.cell.2010.01.018

7 Paul, K., Nieto, V., Carlquist, W. C., Blair, D. F. & Harshey, R. M. The c-di-GMP binding protein YcgR controls flagellar motor direction and speed to affect chemotaxis by a “backstop brake” mechanism. Mol Cell 38, 128–139 (2010). 10.1016/j.molcel.2010.03.001

8 Jenal, U., Reinders, A. & Lori, C. Cyclic di-GMP: second messenger extraordinaire. Nat Rev Microbiol 15, 271–284 (2017). 10.1038/nrmicro.2016.190

9 Leake, M. C. et al. Stoichiometry and turnover in single, functioning membrane protein complexes. Nature 443, 355–358 (2006). 10.1038/nature05135

10 Fukuoka, H., Wada, T., Kojima, S., Ishijima, A. & Homma, M. Sodium-dependent dynamic assembly of membrane complexes in sodium-driven flagellar motors. Mol Microbiol 71, 825–835 (2009). 10.1111/j.1365-2958.2008.06569.x

11 Harman, M. W. et al. The heterogeneous motility of the Lyme disease spirochete in gelatin mimics dissemination through tissue. Proc Natl Acad Sci U S A 109, 3059–3064 (2012). 10.1073/pnas.1114362109

12 Norman, M. U. et al. Molecular mechanisms involved in vascular interactions of the Lyme disease pathogen in a living host. PLoS Pathog 4, e1000169 (2008). 10.1371/journal.ppat.1000169

13 Dunham-Ems, S. M. et al. Live imaging reveals a biphasic mode of dissemination of Borrelia burgdorferi within ticks. J Clin Invest 119, 3652–3665 (2009). 10.1172/JCI39401

14 Chang, Y. et al. Structural insights into flagellar stator-rotor interactions. eLife 8, e48979 (2019). 10.7554/eLife.48979

15 Charon, N. W. et al. The unique paradigm of spirochete motility and chemotaxis. Annu Rev Microbiol 66, 349–370 (2012). 10.1146/annurev-micro-092611-150145

16 Moon, K. H. et al. Spirochetes flagellar collar protein FlbB has astounding effects in orientation of periplasmic flagella, bacterial shape, motility, and assembly of motors in Borrelia burgdorferi. Mol Microbiol 102, 336–348 (2016). 10.1111/mmi.13463

17 Liu, J. et al. Intact flagellar motor of Borrelia burgdorferi revealed by cryo-electron tomography: evidence for stator ring curvature and rotor/C-ring assembly flexion. J Bacteriol 191, 5026–5036 (2009). 10.1128/JB.00340-09

18 Bepler, T., Kelley, K., Noble, A. J. & Berger, B. Topaz-Denoise: general deep denoising models for cryoEM and cryoET. Nat Commun 11, 5208 (2020). 10.1038/s41467-020-18952-1

19 Punjani, A., Rubinstein, J. L., Fleet, D. J. & Brubaker, M. A. cryoSPARC: algorithms for rapid unsupervised cryo-EM structure determination. Nat Methods 14, 290–296 (2017). 10.1038/nmeth.4169

20 Chang, Y. et al. Molecular mechanism for rotational switching of the bacterial flagellar motor. Nature Structural & Molecular Biology 27, 1041–1047 (2020). 10.1038/s41594-020-0497-2

21 Guo, S., Xu, H., Chang, Y., Motaleb, M. A. & Liu, J. FliL ring enhances the function of periplasmic flagella. Proceedings of the National Academy of Sciences 119, e2117245119 (2022). 10.1073/pnas.2117245119

22 Motaleb, M. A., Pitzer Joshua, E., Sultan Syed, Z. & Liu, J. A Novel Gene Inactivation System Reveals Altered Periplasmic Flagellar Orientation in a Borrelia burgdorferi fliL Mutant. Journal of Bacteriology 193, 3324–3331 (2011). 10.1128/jb.00202-11

23 Xu, H., He, J., Liu, J. & Motaleb, M. A. BB0326 is responsible for the formation of periplasmic flagellar collar and assembly of the stator complex in Borrelia burgdorferi. Molecular Microbiology 113, 418–429 (2020). 10.1111/mmi.14428

24 Botting, J. M. et al. FlbB forms a distinctive ring essential for periplasmic flagellar assembly and motility in Borrelia burgdorferi. PLoS Pathog 21, e1012812 (2025). 10.1371/journal.ppat.1012812

25 Santiveri, M. et al. Structure and Function of Stator Units of the Bacterial Flagellar Motor. Cell 183, 244–257.e216 (2020). 10.1016/j.cell.2020.08.016

26 Deme, J. C. et al. Structures of the stator complex that drives rotation of the bacterial flagellum. Nature Microbiology 5, 1553–1564 (2020). 10.1038/s41564-020-0788-8

27 McClune, M. E. et al. The peptidoglycan of Borrelia burgdorferi can persist in discrete tissues and cause systemic responses consistent with chronic illness. Sci Transl Med 17, eadr2955 (2025). 10.1126/scitranslmed.adr2955

28 Chang, Y., Xu, H., Motaleb, M. A. & Liu, J. Characterization of the Flagellar Collar Reveals Structural Plasticity Essential for Spirochete Motility. mBio 12, e0249421 (2021). 10.1128/mBio.02494-21

29 Blair, D. F. Flagellar movement driven by proton translocation. FEBS Lett 545, 86–95 (2003). 10.1016/s0014-5793(03)00397-1

30 Hu, H. et al. Ion selectivity and rotor coupling of the Vibrio flagellar sodium-driven stator unit. Nature Communications 14, 4411 (2023). 10.1038/s41467-023-39899-z

31 Raddi, G. et al. Three-dimensional structures of pathogenic and saprophytic Leptospira species revealed by cryo-electron tomography. J Bacteriol 194, 1299–1306 (2012). 10.1128/JB.06474-11

32 Liu, J. et al. Cellular architecture of Treponema pallidum: novel flagellum, periplasmic cone, and cell envelope as revealed by cryo electron tomography. J Mol Biol 403, 546–561 (2010). 10.1016/j.jmb.2010.09.020

33 Tachiyama, S. et al. FlgY, PflA, and PflB form a spoke-ring network in the high-torque flagellar motor of Helicobacter pylori. Proceedings of the National Academy of Sciences 122, e2421632122 (2025). 10.1073/pnas.2421632122

34 Tachiyama, S. et al. The flagellar motor protein FliL forms a scaffold of circumferentially positioned rings required for stator activation. Proceedings of the National Academy of Sciences 119, e2118401119 (2022). 10.1073/pnas.2118401119

35 Feng, X. et al. Structural insights into the assembly and evolution of a complex bacterial flagellar motor. Nature Microbiology 11, 770–785 (2026). 10.1038/s41564-025-02248-5

36 Beeby, M. et al. Diverse high-torque bacterial flagellar motors assemble wider stator rings using a conserved protein scaffold. Proc Natl Acad Sci U S A 113, E1917–1926 (2016). 10.1073/pnas.1518952113

37 Drobnic, T. et al. In situ structure of a bacterial flagellar motor at subnanometre resolution reveals adaptations for increased torque. Nat Microbiol 10, 1723–1740 (2025). 10.1038/s41564-025-02012-9

38 Mastronarde, D. N. Automated electron microscope tomography using robust prediction of specimen movements. J Struct Biol 152, 36–51 (2005). 10.1016/j.jsb.2005.07.007

39 Zheng, S. Q. et al. MotionCor2: anisotropic correction of beam-induced motion for improved cryo-electron microscopy. Nat Methods 14, 331–332 (2017). 10.1038/nmeth.4193

40 Bepler, T. et al. Positive-unlabeled convolutional neural networks for particle picking in cryo-electron micrographs. Nat Methods 16, 1153–1160 (2019). 10.1038/s41592-019-0575-8

41 Sanchez-Garcia, R. et al. DeepEMhancer: a deep learning solution for cryo-EM volume post-processing. Communications Biology 4, 874 (2021). 10.1038/s42003-021-02399-1

42 Cao, H. et al. EMReady2: improvement of cryo-EM and cryo-ET maps by local quality-aware deep learning with Mamba. Nature Communications 17, 6493 (2026). 10.1038/s41467-026-71794-1

43 Rego, R. O., Bestor, A. & Rosa, P. A. Defining the plasmid-borne restriction-modification systems of the Lyme disease spirochete Borrelia burgdorferi. J Bacteriol 193, 1161–1171 (2011). 10.1128/JB.01176-10

44 Li, C. et al. Asymmetrical flagellar rotation in Borrelia burgdorferi nonchemotactic mutants. Proc Natl Acad Sci U S A 99, 6169–6174 (2002). 10.1073/pnas.092010499

45 Zhang, K. et al. FlhF regulates the number and configuration of periplasmic flagella in Borrelia burgdorferi. Mol Microbiol 113, 1122–1139 (2020). 10.1111/mmi.14482

46 Sze, C. W. et al. Study of the response regulator Rrp1 reveals its regulatory role in chitobiose utilization and virulence of Borrelia burgdorferi. Infection and immunity 81, 1775–1787 (2013). 10.1128/IAI.00050-13

47 Sze, C. W. et al. The Flagellin-Specific Chaperone FliS of Borrelia burgdorferi Controls the Cytoplasmic Pool of Flagellins at the Level of Translation Initiation, Secretion, and Proteolysis. Mol Microbiol 124, 173–187 (2025). 10.1111/mmi.15380

48 Zhang, K., Sze, C. W., Zhao, H., Liu, J. & Li, C. Borrelia burgdorferi serine protease HtrA is a pleiotropic regulator of stress response, motility, flagellar hemostasis, and infectivity. Commun Biol 8, 341 (2025). 10.1038/s42003-025-07781-x

49 Bakker, R. G., Li, C., Miller, M. R., Cunningham, C. & Charon, N. W. Identification of specific chemoattractants and genetic complementation of a Borrelia burgdorferi chemotaxis mutant: flow cytometry-based capillary tube chemotaxis assay. Appl Environ Microbiol 73, 1180–1188 (2007). 10.1128/AEM.01913-06

50 Zhang, K. et al. Two CheW coupling proteins are essential in a chemosensory pathway of Borrelia burgdorferi. Mol Microbiol 85, 782–794 (2012). 10.1111/j.1365-2958.2012.08139.x

51 Motaleb, M. A. et al. Borrelia burgdorferi periplasmic flagella have both skeletal and motility functions. Proc Natl Acad Sci U S A 97, 10899–10904 (2000). 10.1073/pnas.200221797

52 Xu, A. & Xu, C. FastTomo: A SerialEM Script for Collecting Electron Tomography Data. bioRxiv (2021). 10.1101/2021.03.16.435675

53 Kremer, J. R., Mastronarde, D. N. & McIntosh, J. R. Computer visualization of three-dimensional image data using IMOD. J Struct Biol 116, 71–76 (1996). 10.1006/jsbi.1996.0013

54 Agulleiro, J. I. & Fernandez, J. J. Tomo3D 2.0--exploitation of advanced vector extensions (AVX) for 3D reconstruction. J Struct Biol 189, 147–152 (2015). 10.1016/j.jsb.2014.11.009

55 Winkler, H. 3D reconstruction and processing of volumetric data in cryo-electron tomography. J Struct Biol 157, 126–137 (2007). 10.1016/j.jsb.2006.07.014

56 Winkler, H. et al. Tomographic subvolume alignment and subvolume classification applied to myosin V and SIV envelope spikes. Journal of Structural Biology 165, 64–77 (2009). 10.1016/j.jsb.2008.10.004

57 Abramson, J. et al. Accurate structure prediction of biomolecular interactions with AlphaFold 3. Nature 630, 493–500 (2024). 10.1038/s41586-024-07487-w

58 Goddard, T. D. et al. UCSF ChimeraX: Meeting modern challenges in visualization and analysis. Protein Sci 27, 14–25 (2018). 10.1002/pro.3235

59 Emsley, P. & Cowtan, K. Coot: model-building tools for molecular graphics. Acta Crystallogr D Biol Crystallogr 60, 2126–2132 (2004). 10.1107/S0907444904019158

60 Liebschner, D. et al. Macromolecular structure determination using X-rays, neutrons and electrons: recent developments in Phenix. Acta Crystallogr D Struct Biol 75, 861–877 (2019). 10.1107/S2059798319011471

61 Sievers, F. & Higgins, D. G. Clustal Omega for making accurate alignments of many protein sequences. Protein Sci 27, 135–145 (2018). 10.1002/pro.3290

